# Screening Molecular Recognition Element-Based SWCNT Optical Sensors for the Inflammatory Cytokine TNF-α

**DOI:** 10.64898/2026.01.22.701124

**Authors:** Syeda Rahman, Atara R. Israel, Amelia Ryan, Ryan Williams

**Author notes:** these authors contributed equally to this work (Syeda Rahman, Atara R. Israel, Amelia Ryan).

## Abstract

TNF-α (Tumor Necrosis Factor) is a proinflammatory cytokine that amplifies inflammatory response and promotes leukocyte recruitment. TNF-α is primarily produced by activated macrophages, among others, in response to infection, inflammation, or tissue damage. Given its central role in normal and abnormal immune responses, it is the target of several therapeutics, such as adalimumab and etanercept. TNF-α is also a prognostic and diagnostic biomarker associated with Rheumatoid Arthritis, Alzheimer’s disease, Multiple Sclerosis, several kidney diseases, several cancers, Type 2 diabetes, sepsis, and others. Spatial quantification of TNF-α in disease models can also be a powerful tool to understand the contributions of inflammatory processes to disease progression. Single-walled carbon nanotubes (SWCNT) are cylindrical carbon lattices that emit distinct near-infrared bandgap photoluminescence. In this work, we evaluated three aptamer-based sensor constructs, plus an additional two iterations of one aptamer sequence, and two antibody-based sensor constructs for TNF-α that use SWCNT near-infrared photoluminescence signal transduction. Several, but not all, of these aptamer and antibody-based sensors sensitively and selectively detected TNF-α in serum in a physiologically relevant range, and we found that their sensing was improved by both passivation and incorporating an exogenous quencher onto the aptamer sequence. Specifically, we found that modification of one aptamer sequence with a Black Hole Quencher induced selective detection in serum when passivated with poly-L-Lysine. This study highlights the importance, and challenges, of translating previously-validated molecular recognition elements to new detection conditions, in this case on the surface of SWCNT and in challenging serum conditions. It also validated a lead sensor construct that builds upon constructs that failed in serum. We anticipate that the sensors evaluated here will have utility in both the diagnosis and study of inflammation-driven chronic disease, while the sensor assessment framework will help drive the broader field of molecularly specific diagnostics.

## Introduction

Tumor Necrosis Factor α (TNF-α) is a pleiotropic cytokine which has regulatory effects on the body’s inflammatory response and is involved in the pathogenesis of many inflammation-linked diseases.^1-3^ This homotrimer protein is mainly produced by activated macrophages, T-lymphocytes and natural killer cells, and it can trigger other cytokines and chemokines to upregulate inflammatory response.^4^ Although TNF-α is a crucial signaling molecule at normal levels, excess secretion is a key factor in disease pathophysiology. Dysregulated TNF-α expression can contribute to autoimmune diseases such as rheumatoid arthritis, inflammatory bowel disease, psoriatic arthritis, psoriasis, among other conditions.^4^ Therefore, quantitative detection of TNF-α holds value for disease diagnosis and in the study of inflammatory disease onset and progression. Standard methods for detecting TNF-α include mass spectrometry and immunoassays, though these are costly, time-consuming, and require expert user operation. In addition, these are destructive methods that are used *ex vivo*. In order to have a better understanding of TNF-α temporal and spatial disease contributions, it is necessary to develop alternative detection methods that exhibit rapid response time, low cost, *in situ* readout, and ease of use.

Single-walled carbon nanotubes (SWCNT) consist of a sp^2^-hybridized carbon lattice structure that can be conceptualized as a single sheet of rolled graphene. They are quasi-one-dimensional nanoparticles with a diameter of 0.5 to 2 nm and length of up to 1 mm. The carbon lattice structure of SWCNT is defined by a chiral index, denoted with (*n,m*) coordinates, which determines its optoelectrical properties.^5^ SWCNT exhibit near-infrared (NIR) photoluminescence across the optical bandgap, and each (*n, m*) species absorbs and emits at distinct wavelengths. Near-infrared SWCNT fluorescence is particularly useful for biosensor transduction as it does not photobleach and exhibits substantial tissue penetration depth with minimal autofluorescence in biological tissues.^5, 6^ SWCNTs are extremely sensitive to their local dielectric environment, which can be directed via corona phase interactions or functionalization with a molecular recognition probe.^5^ Surface functionalization of SWCNTs can aid in their selectivity and specificity towards biological targets and is necessary to solubilize them. Aptamers, synthetic oligonucleotides with binding affinity for a specific target, have been used to functionalize SWCNT – increasing biocompatibility and acting as a recognition probe for the target molecules. Our previous studies have demonstrated SWCNT nanosensor functionalized with ssDNA aptamers for the detection of interleukin-6 and cortisol.^7, 8^ We have also conjugated antibodies to ssDNA-SWCNT.^9, 10 11, 12^

In this work, we assessed the sensitivity, selectivity, and robustness of several sensor constructs designed to detect TNF-α using a SWCNT optical transducer. In our experience with incorporating both aptamers and antibodies into SWCNT optical sensors, we typically find that it is necessary to test several constructs prior to actually using the sensor. Here, we evaluated three separate TNF-α specific aptamer sequences, plus two additional variations of one sequence, and two TNF-α antibodies as recognition elements for SWCNT-based optical sensors. We also explored thermal and ion-induced conformational changes in the aptamers to enhance sensor response. We evaluated the selectivity of our sensors in the presence of competing serum proteins with and without passivation agents to block nonspecific protein adsorption to the SWCNT surface. This work represents a rational framework for sensor design, screening, and optimization that may be broadly applicable to a wide variety of analytes.

## Methods

### SWCNT suspension with ssDNA

HiPCO single walled carbon nanotubes (SWCNT) (NanoIntegris Technologies, Boisbriand, Quebec) were suspended in solution separately with seven oligonucleotide sequences, described below (**Table 1**; Integrated DNA Technologies, Coralville, IA), at a 1:2 SWCNT:ssDNA mass ratio in 1X PBS as previously described.^13^ The sample was sonicated at 40% amplitude for 60 minutes in an ice bath. The resulting suspension was ultracentrifuged at 58000 x g for 1 hour (Beckman Coulter; California, USA) to remove impurities and aggregates. The top 75% of the centrifuged suspension was collected and stored for use. Immediately prior to use, SWCNT suspensions were filtered with a 100 kDa centrifugal filter (Millipore Sigma, Burlington, MA) to remove excess unbound oligonucleotides. The solution was then resuspended in 100-200 µl of 1X PBS.

**Table 1:**
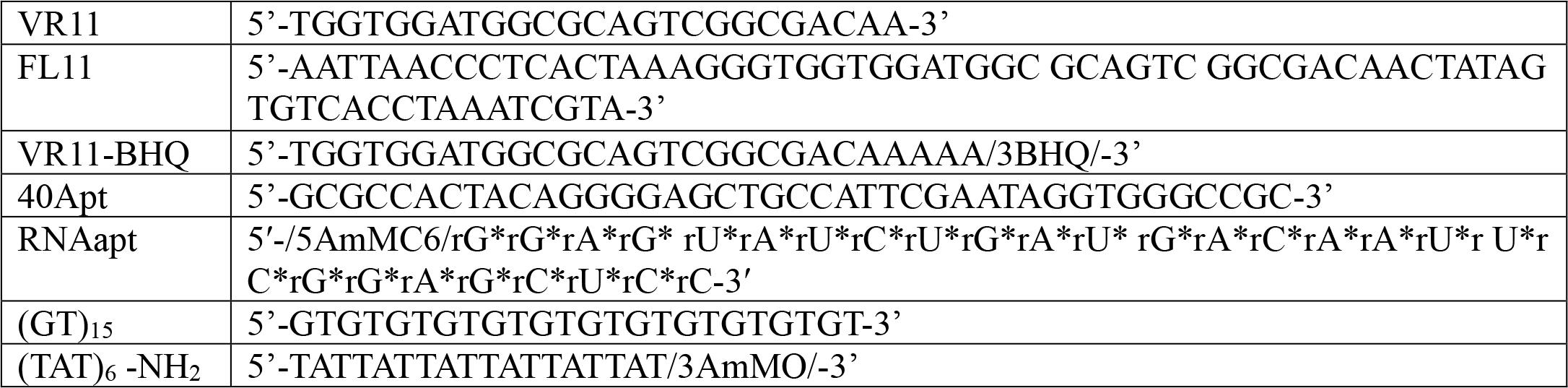
Oligonucleotide sequences used to suspend SWCNT.

#### Oligonucleotides used for SWCNT suspension

##### VR11

Previously published as Variable Region 11 (VR11), in vitro selection was performed to obtain an aptamer to recognize TNF-α and block its activity *in vitro*. It demonstrated a dissociation constant of 7.0 ± 2.1 nM and inhibited apoptosis induced by TNF-α and production of nitric oxide.^14^

##### FL11

This sequence (Full Length 11) is a longer version of VR11. It inhibited TNF-α functionality *in vitro*.^14^ This longer sequence has a higher G-rich content, potentially imparting more structural stability and binding to TNF-α.

##### VR11-BHQ

We modified VR11 with a quencher dye BHQ (Black Hole Quencher) and spacer sequence (five adenines) to enhance the response of VR11 in serum.^15-20^ We hypothesized that potential aptamer conformational changes upon binding would induce a modification of SWCNT fluorescence due to the proximity of BHQ to the nanotube.

##### 40Apt

In published studies, this aptamer demonstrated a strong binding affinity for TNF-α and blocked its functions in Acute Lung Injury and Acute Liver Failure *in vivo* with a dissociation constant of 8 nM against human TNF-α.^21^

##### RNAapt

This RNA aptamer was previously selected and demonstrated to bind to TNF-α. It was subsequently deployed in an electrochemical sensor, with functionality in whole blood.^22^

##### (GT)_15_

This control sequence was chosen as it is known to stably suspend SWCNT and it has no particular selective affinity for TNF-α protein, though it does allow for responses to other proteins and analytes.^23-27^ It was used as a control sequence to evaluate the specific binding nature of TNF-α molecular recognition elements compared to that of ssDNA in general.

##### (TAT)_6_-NH_2_

This functionalized oligonucleotide sequence has been used in prior studies to stably encapsulate SWCNT and allow for carbodiimide conjugation chemistry to an antibody.^9, 11, 28^

### SWCNT absorbance characterization

SWCNT suspended by oligonucleotides were characterized with a V-730 UV-Visible absorption spectrophotometer measured over 300-1100 nm (Jasco Inc., Easton, MD). We determined suspension concentration using the molar extinction coefficient Abs_630_= 0.036 L mg−1 cm−1 as previously described.^7^

### Antibody conjugation to ssDNA-SWCNT

We used SWCNT dispersed with (TAT)_6_-NH_2_ to covalently conjugate a monoclonal antibody (mAb) (Invitrogen, Waltham, MA, RRID: AB_468487) and a polyclonal antibody (pAb) (Invitrogen, Waltham, MA, RRID: AB_2609680) using carbodiimide crosslinking as previously described.^9, 11, 28^ The carboxyl groups of the antibody were activated with 25x molar excess of N-hydroxysuccinimide (NHS) (TCI Chemicals, Portland, OR) and 10x excess of 1-ethyl-3-(3-dimethylainopropyl)carbodiimide (EDC) (Sigma Aldrich, St. Louis, MO) for 15 minutes at 4°C. The reaction was quenched with 1 µl of 2-mercaptoethanol (Sigma Aldrich, St. Louis, MO). The SWCNT-(TAT)_6_-NH_2_ dispersion was added to the activated antibody in an equimolar ratio and incubated at 4°C for 2 hours with gentle vortex agitation every 30 minutes. The sample was dialyzed against deionized water for 48 hours to remove excess reagents in a 1000 kDa molecular weight cutoff filter (Spectrum Labs, Rancho Dominguez, California) with three dialysate exchanges. Dynamic Light Scattering (DLS) and ζ-potential were performed to estimate the size and charge, respectively, to confirm successful conjugation (Malvern ZS-90, Westborough, MA).

### SWCNT-Ab and SWCNT-aptamer surface passivation

Surface passivation can enhance the response of SWCNT sensors by inhibiting nonspecific protein adsorption on the nanotube surface and directing analyte interactions to the molecular recognition element. After screening all constructs for their responsiveness to TNF-α, two passivation agents were explored to maximize the response of the sensors. Bovine serum albumin (BSA; Fisher, Waltham, MA) and poly-L-lysine (PLK; Advanced Biomatrix, Carlsbad, CA) have previously been identified as effective passivation agents for SWCNT optical sensors.^9, 11, 28^ For experiments in which sensor function under passivation conditions were tested, functionalized SWCNTs were incubated with a 50x mass excess of a given passivation agent at 4°C for 30 minutes prior to sensor deployment.

### Near-infrared fluorescence spectroscopy

Near-infrared fluorescence spectra of SWCNT sensors were acquired via a ClaIR custom-built NIR plate reader (Photon etc., Montreal, Quebec) with laser source excitation wavelengths 655 nm and 730 nm. Near-infrared spectral acquisitions were performed in a 96-well plate. Spectra were acquired between 900 and 1700 nm with excitation laser power of 1750 mW and an exposure time of 500 ms.

### Screening sensor responses to TNF-α

To evaluate the response of all sensor constructs to TNF-α, we diluted each sensor to 1 mg/L in 1X PBS in a total volume of 120 µl. Each construct was assessed in triplicate. An initial baseline measurement NIR fluorescence was obtained, then recombinant TNF-α (R&D Systems, Minneapolis, MN) was added across a range of concentrations from 1-250 nM for the experimental groups. Control untreated samples received an equal volume of 1x PBS only. NIR fluorescence measurements were acquired every 15 minutes for 2.5 hours to evaluate fluorescence modulation over time. We then challenged each sensor with heat-inactivated fetal bovine serum (FBS; Corning Inc., Corning, NY) to simulate complex biological conditions. Sensor samples were prepared as described above with the addition of 10% FBS. TNF-α protein was then added to the sensors and measurements were acquired as previously described.

### Temperature and divalent metal ion-induced aptamer conformational folding

Generally, aptamers adopt a three-dimensional structure that allows them to bind to their target. When the aptamer sequences were introduced to SWCNT during sonication, they may exist in non-optimal conformations on the surface of the nanotube. We sought to assess aptamer-based sensor function in conditions which would promote optimal aptamer folding after sonication and purification. We did so by heat-denaturing the aptamer and introducing divalent metal ions to sequentially unfold and refold the aptamer into an optimal conformation. Thermal denaturation and refolding of each aptamers was achieved by heating the aptamer-SWCNT complex at 95°C for 5 minutes and letting it cool at room temperature for 20 minutes, then adding 1mM of MgCl_2_ before obtaining fluorescence measurements as above.^29^

### Assessment of sensor selectivity

To further test the selectivity of the VR11, FL11, and VR11-BHQ sensors to TNF-α, equal concentrations of bovine serum albumin (BSA; Fisher, Waltham, MA), interleukin-1β (IL-1β; Peprotech, Cranbury, NJ) and interleukin-6 (IL-6; Gibco, Waltham, MA) were added to the sensor similarly to TNF-α sensitivity experiments. Their response was evaluated over 2.5 hours every 15 minutes.

### Data processing and analysis

All experiments were performed in triplicate. Baseline measurements were acquired before test protein addition to benchmark change in sensor center wavelength and emission intensity. Individual SWCNT (*n,m*) emission peaks were identified according to published studies.^30, 31^ Each peak was fit using a pseudo-Voight model with a custom MATLAB code (code available upon request). Center wavelength and intensity measurements were used in analyses only when model fit R^2^ were greater than 0.95. Triplicate averages and propagated standard deviations were obtained and reported. Statistical significance was determined with a two-sample *t* test.

## Results and Discussion

We designed seven SWCNT-based optical sensors to detect TNF-α. The sensor complexes were synthesized by encapsulating HiPCO-produced SWCNT with oligonucleotide sequences (**Table 1**), including TNF-α aptamers and a non-specific sequence as an intermediate linker for subsequent antibody conjugation. Optical characterization of the SWCNT suspensions (**Figure 1, Supplementary Figure 1).^11, 12, 32, 33^**

**Figure 1.**
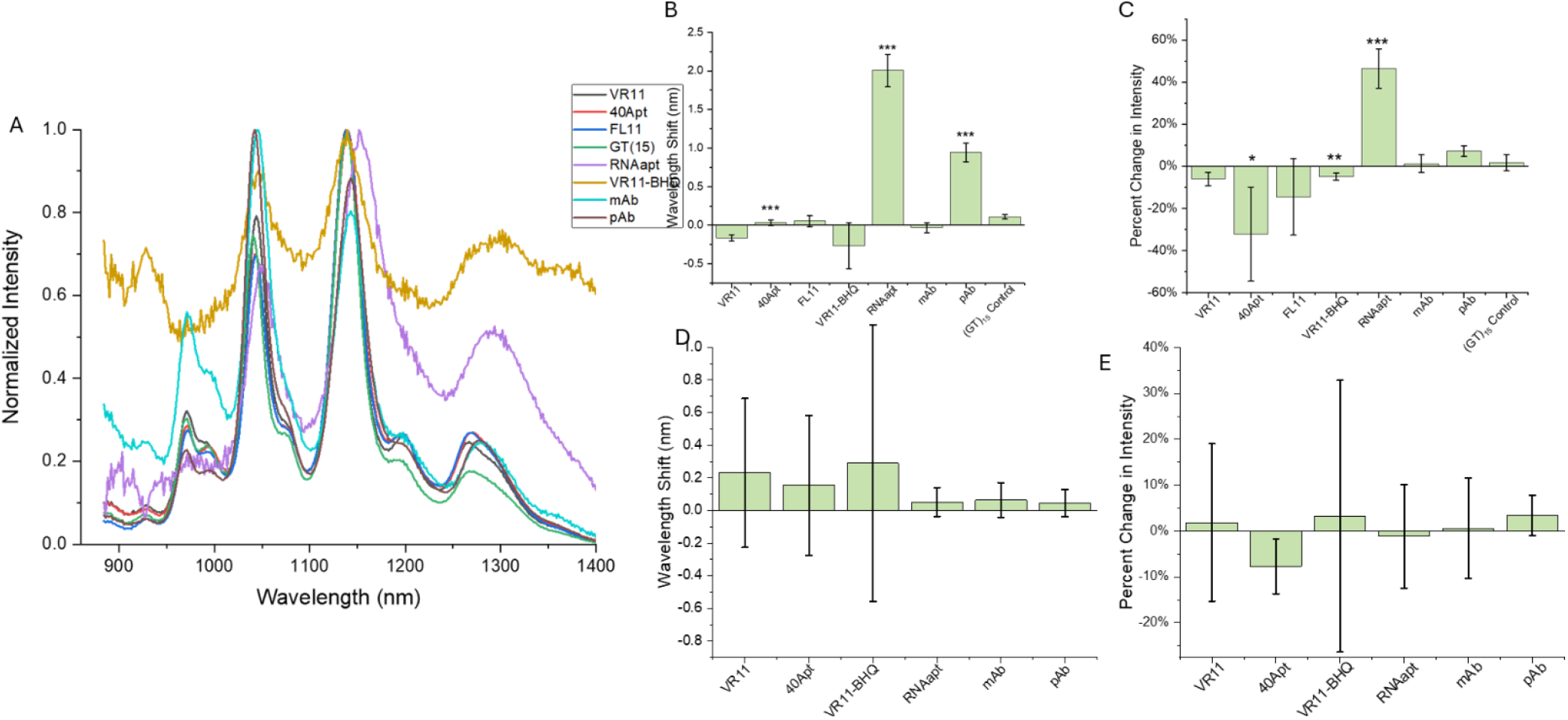
TNF-α sensor screening. A) NIR fluorescence spectra of all SWCNT constructs following 655 nm excitation. B) Center wavelength shift of the (7,5) SWCNT for all sensor constructs after three hours of incubation with 250 nM TNF-α protein in 1X PBS. C) Change in (7,5) emission intensity after three hours of incubation with 250 nM TNF-α protein in 1X PBS. D) Center wavelength shift of the (7,5) SWCNT for each sensor construct after three hours of incubation with 250 nM TNF-α protein in 10% FBS. E) Change in (7,5) emission intensity after three hours of incubation with 250 nM TNF-α protein in 10% FBS.

**Figure 1**) determined that most were well-dispersed and exhibited bright near-infrared fluorescence. The fluorescence of the VR11-BHQ was somewhat dim due to either presence of the quencher or overall efficiency of suspension. However, it had identifiable and evaluable (*n,m*) peaks. Similarly, the RNAapt-SWCNT sensor exhibited less defined peaks than ssDNA-SWCNT, though prominent peaks were clear. It is interesting to observe a slight red-shift in the RNAapt-SWCNT sensor baseline fluorescence compared to others. It is equally interesting that we observed a smaller redshift for the antibody-conjugated SWCNT compared to the others. Further characterization was performed on the SWCNT-Ab formulations, wherein DLS found that the SWCNT complexes were larger in size after the conjugation process, as expected from our prior work.^11, 12, 32, 33^ Zeta potential measurements indicated that the surface charge of the constructs increased after antibody conjugation, suggesting successful conjugation of the antibodies to the nanotube complexes (**Supplementary Figure 1**).^11, 12, 32, 33^

### Comparative assessment of molecularly-specific TNF-α sensors

We primarily analyzed the (7,5) SWCNT species (**Figure 1**), though similar results were found for the (7,6) and (9,4) SWCNT species (**Supplementary Figure 2)**. SWCNT functionalized with 40Apt, RNAapt, and pAb exhibited a statistically significant red shift in response to TNF-α, with an average shift of 0.03 nm, 2.0 nm, and 0.9 nm, respectively (**Figure 1B**). Though the magnitude of the shift seen in 40Apt is exceedingly small and not suitable for sensor development, it was repeatable. The RNAapt and pAb shifts are, however, both robust and significant. We also observed significant changes in fluorescence intensity, specifically 40Apt, VR11-BHQ, and RNAapt (**Figure 1C**).

Almost all SWCNT sensor constructs demonstrated a significant change in either wavelength shift, intensity, or both, in response to TNF-α in buffer conditions for at least one of the (*n,m*) species analyzed. The mAb-functionalized sensor did not exhibit a significant response, though our prior experience is that this is likely a product of the antibody compatibility with the assay, rather than the platform. RNAapt emerged as a particularly strong candidate as it enabled a wavelength shift across (7,5), (7,6), and (9,4) SWCNT and intensity modulation in the (7,5) and (7,6). SWCNT functionalized with pAb also exhibited wavelength shifts for two out of the three (*n,m*) analyzed. The control SWCNT-(GT)_15_ did show a statistically significant, but very small in magnitude, shift and change in intensity of the (7,6) peak, but no change in the other two SWCNT species assessed.

The VR11 family of sensors was particularly interesting due to the various iterations assessed. VR11 enabled a wavelength-based response for the (7,5) and (9,4) SWCNT. The quencher-attached iteration, VR11-BHQ, exhibited a wavelength shift for the (7,6) SWCNT and a minimal fluorescence intensity modulation for all SWCNT species analyzed. This indicates some potential quencher-induced charge transfer but not modification of the local dielectric – perhaps the quencher serves as an anchor on the SWCNT surface. The longer, full-length version of VR11 (FL11), enabled a small shift in (7,6) SWCNT fluorescence, however its overall response to TNF-α was less robust than VR11. We posit that FL11 exhibits less conformational rearrangement on the SWCNT surface upon analyte binding compared to VR11.

Functional sensor response was then assessed in the presence of 10% serum to simulate a protein-rich biological environment. However, none of the sensors exhibited a significant response to 250 nM TNF-α for any of the species analyzed (**Figures 1D-E, Supplementary Figure 2)**. This is not uncommon, as it is well-known that ionic conditions and proteins in serum can interfere with binding and specific signaling. Thus, we further investigated the mechanisms of sensor response for each of these sensor constructs to understand how to improve sensor functionality.

### In-depth assessment of each sensor construct

#### Control (GT)_15_, not specific for TNF-α

The SWCNT-(GT)_15_ complex was tested in buffer conditions with the addition of 100 nM and 250 nM TNF-α protein. In buffer conditions, SWCNT-(GT)_15_ exhibited negligible wavelength shifts in response to TNF-α compared to controls with no protein added (**Supplementary Figure 3**). The only significant difference we observed was a <0.2 nm change for 250 nM TNF-α compared to PBS. We also observed no changes in fluorescence intensity in response to TNF-α.

*VR11 aptamer family*: The VR11 aptamer was incubated with 1 – 500 nM TNF-α. The results of the (7,5) SWCNT (**Figures 2A-B**) demonstrate a small-in-magnitude, but statistically significant, detection of TNF-α in comparison to a PBS control at all concentrations except for 2.5 nM. TNF-α responses were also analyzed for the (7,6) and (9,4) SWCNT (**Supplementary Figure 4**), although it should be noted here that sensor responses, specifically to 250 nM TNF-α, were not reproducible. Since the VR11 sequence has previously been assessed and validated for the detection of TNF-α, we decided to conduct further testing of the sensor construct to determine its specificity and selectivity.^34, 35^ We challenged the sensor with equal concentrations of cytokines interleukin 1 β (IL-1β) and interleukin 6 (IL-6). Neither the (7,5) nor the (7,6) exhibited significant intensity responses, however it is interesting that VR11-SWCNT did exhibit a small, but significant, blue shift in response to TNF-α and a red shift in the presence of IL-1β and IL-6 (**Figures 2C-D, Supplementary Figure 4**). We also compared the influence of thermal denaturing the aptamer-functionalized SWCNT to see if heat-induced conformational change influences the response to TNF-α. However, despite some wavelength response, substantial variability existed, possibly due to instability of the ssDNA-SWCNT construct under high heat (**Figures 2E-F)**.

**Figure 2.**
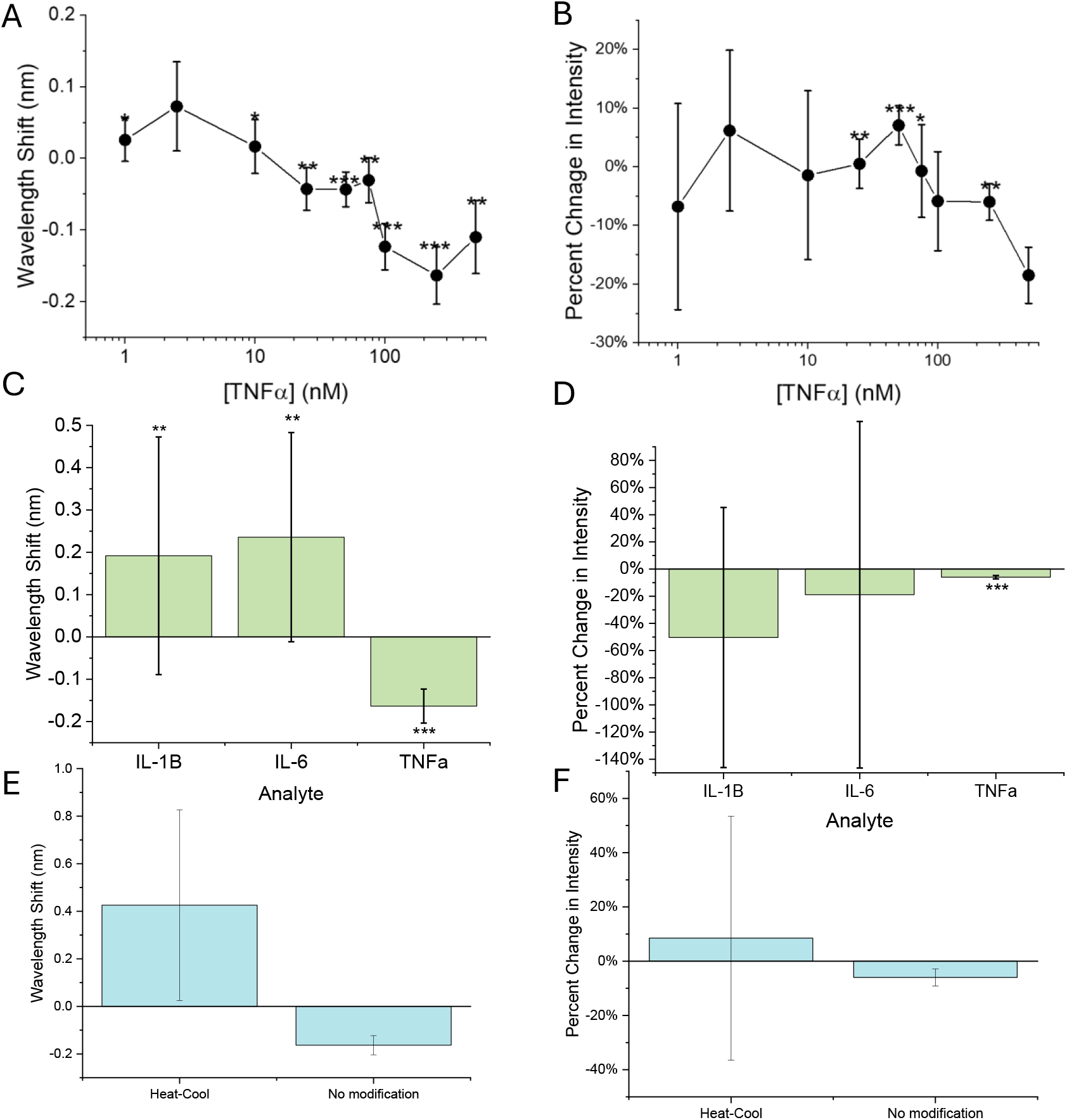
VR11-SWCNT detection of TNF-α. A) Wavelength shift of the (7,5) as a function of TNF-α concentration after 180 minutes of incubation in 1x PBS. B) Changes in (7,5) emission intensity as a function of TNF-α concentration after 180 minutes of incubation in 1x PBS. C) Wavelength shift of the (7,5) SWCNT after incubation with 250 nM IL-1β, IL-6, and TNF-α. D) Modulation of (7,5) emission intensity after 180 minutes incubation with 250 nM IL-1β, IL-6, and TNF-α. E) Wavelength shift of (7,6) SWCNT after a 180 minute incubation with 100 nM TNF-α following heat-cool. F) Modulation of the (7,6) emission intensity after 180 minute incubation with TNF-α following heat-cool. Mean represents average of triplicate. Error bars represent +/-standard deviation. T-test significance indicated by * =p<0.05, **=p<0.01, ***=p<0.001.

In an effort to enhance the response of the nanosensor to TNF-α, we evaluated a full-length variant, FL11, as the addition of flanking sequences may improve DNA-SWCNT sensor response.^14, 36, 37^ We found that FL11 did not exhibit a substantial response to TNF-α. To determine its selectivity towards TNF-α, we challenged the FL11-SWCNT construct with both TNF-α and BSA at 250 nM. The nanosensor exhibited a wavelength shift and intensity changes in the presence of BSA as well as the presence of TNF-α, (**Supplementary Figure 5**), indicating lack of specificity towards TNF-α. It is possible that the conformation of the aptamer was disrupted during the functionalization process onto the SWCNT surface.

To further amplify the VR11-SWCNT sensor response, we added a Black Hole Quencher (BHQ) to the aptamer. We hypothesized that in the absence of TNF-α, the location of the quencher would be closer to the surface of the nanotube, causing its baseline fluorescence to be quenched. However, in the presence of TNF-α, the analyte would bind to the aptamer, decouple the BHQ from the SWCNT surface, and restore fluorescence intensity.^38^ This phenomenon was somewhat evident during the screening process, noting significant intensity changes of the nanosensor in the presence of 250 nM TNF-α in buffer conditions for the (7,5) and (9,4) species and a wavelength shift in the (7,6) species (**Figure 1C, Supplementary Figure 2A, F**). This response, however, was not seen when the nanosensor was tested in a more complex environment (**Figures 1D-E**). In an experiment to compare response to TNF-α and BSA at equal concentrations, the nanosensor exhibited a significant, substantial 2.5 nm wavelength shift for the (7,6) (**Figure 3A**) and of 1.4 nm for the (9,4) when incubated with TNF-α (**Supplementary Figure 6**). We also attempted to passivate the sensor using poly-L-lysine (PLK) to improve its response in more complex environments. This passivated (7,6) SWCNT demonstrated a significant wavelength shift in comparison to its non-passivated (7,6) in the presence of 500 nM TNF-α. Similar responses were seen in the (7,5) and (9,4) SWCNT as well (**Figure 3C, Supplementary Figure 6)**. Despite this, there was negligible change in fluorescence intensity after TNF-α incubation with passivated VR11-BHQ-SWCNT (**Figure 3D**).

**Figure 3.**
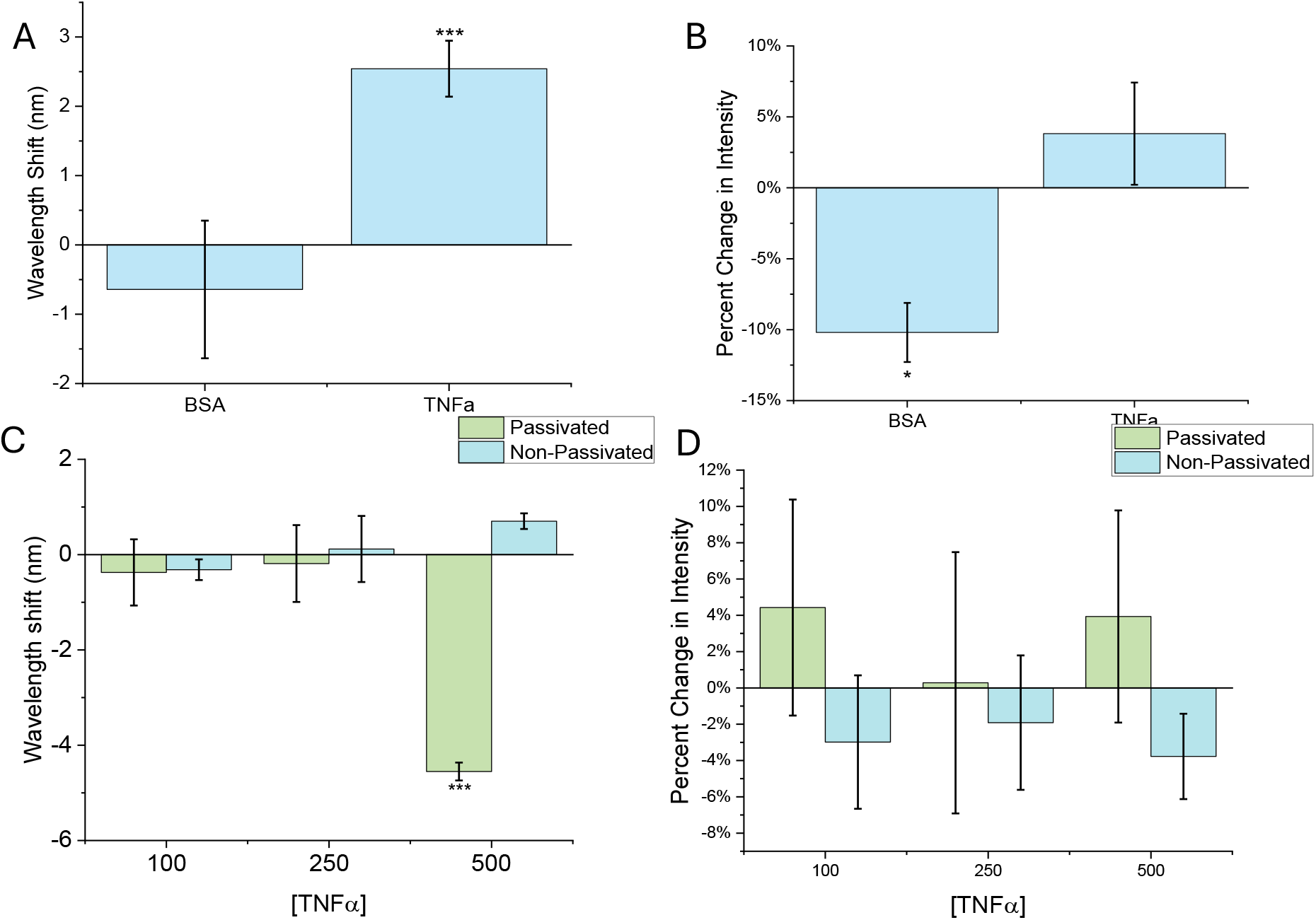
Assessment of VR11-BHQ-SWCNT specificity to TNF-α. A) Wavelength shift of the (7,6) SWCNT in the presence of 250 nM BSA and TNF-α after 180 minutes in 1x PBS. B) Change in (7,6) emission intensity in the presence of 250 nM BSA and TNF-α after 180 minutes in 1x PBS. C) Wavelength shift of the (7,6) SWCNT after 180 minutes in the presence of TNF-α in 10% FBS with and without PLK passivation. D) Intensity change in the (7,6) after 180 minutes in the presence of TNF-α in 10% FBS with and without PLK passivation. Mean represents average of triplicate. Error bars represent +/-standard deviation. T-test significance indicated by ***=p<0.001.

#### 40-Base ssDNA aptamer (40Apt)

40Apt is a ssDNA aptamer, separate from VR11, that was selected for its affinity for TNF-α.^21^ We evaluated sensitivity of SWCNT-40Apt by first screening it against 250 nM TNF-α in buffer conditions. We found small but significant wavelength shift responses in all (*n,m*) species analyzed and significant intensity quenching in two species. In 10% FBS, we did not observe substantial responses to TNF-α (**Figures 1B-E**). We then assessed sensor response across a broader concentration range (**Figures 4A-B**), finding minimal wavelength shifts for all chiralities analyzed with no clear response pattern (**Supplementary Figure 7**). We next added 1 mM MgCl_2_ to 40Apt-SWCNT to induce adoption of appropriate three-dimensional aptamer conformation.^29^ Separately, the nanosensor

**Figure 4.**
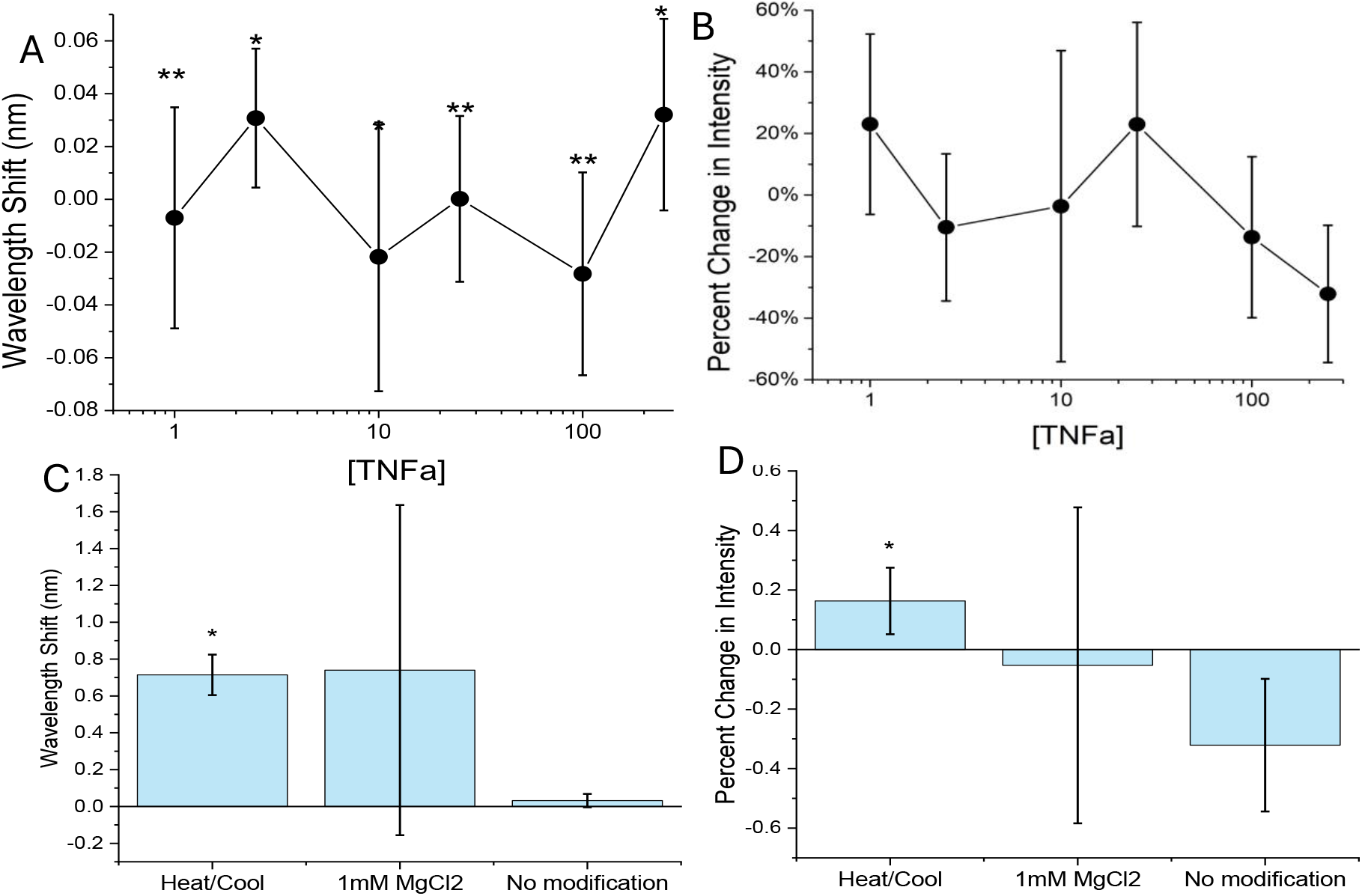
40Apt-SWCNT response to TNF-α. A) Wavelength shift of the (7,5) SWCNT as a function of TNF-α concentration after 180 minutes of incubation in 1x PBS. B) Change in (7,5) emission intensity after 180 minutes of incubation. C) Wavelength shift and D) intensity changes after 180 minutes incubation following heat-cool and divalent cation refolding of 40Apt-SWCNT. Mean represents average of triplicate. Error bars represent +/-standard deviation. T-test significance indicated by *=p<0.05.

also underwent thermal denaturation and subsequent cooling to facilitate conformation adoption.^39^ Interestingly, the heat-cool process enhanced sensor response to 250 nM TNF-α, facilitating a 0.7 nm wavelength shift, whereas divalent ion addition appeared to induce substantial variability (**Figures 4C-D**).

#### ssRNA aptamer for TNF-α (RNAapt)

Some studies have demonstrated that RNA aptamers have enhanced binding affinity compared to DNA aptamers.^40^ We evaluated a TNF-α-specific RNA aptamer SWCNT sensor. Upon initial screening with 250nM TNF-α **(Figure 1)**, we saw a substantial wavelength shift across all (*n,m*) analyzed, along with significant intensity differences in two out of three species analyzed. In serum conditions, SWCNT-RNAapt response TNF-α was negligible without passivation (**Figures 5C-D**). To attempt to improve response in serum, the sensor was passivated using PLK and deployed, though there remained substantial variability (**Figures 5C-D**). The sensor was then tested against equal concentrations of BSA and TNF-α to test its specificity. We found a 2-fold greater shift and change in intensity modulation for each (*n,m*) species (**Figures 5E-F; Supplementary Figure 8**) in response to TNF-α compared to BSA. This aptamer has not been widely studied in this context, and few RNA aptamer have been assessed with SWCNT. It does, however, hold promise for development into a specific and selective TNF-α nanosensor with improved passivation schemes in serum. However, literature on SWCNT-RNA interactions has shown that time-dependent fluorescence variability is higher in these hybrids than in SWCNT-DNA constructs, therefore the sensor’s long-term fluorescence stability should be further evaluated.^23^

**Figure 5.**
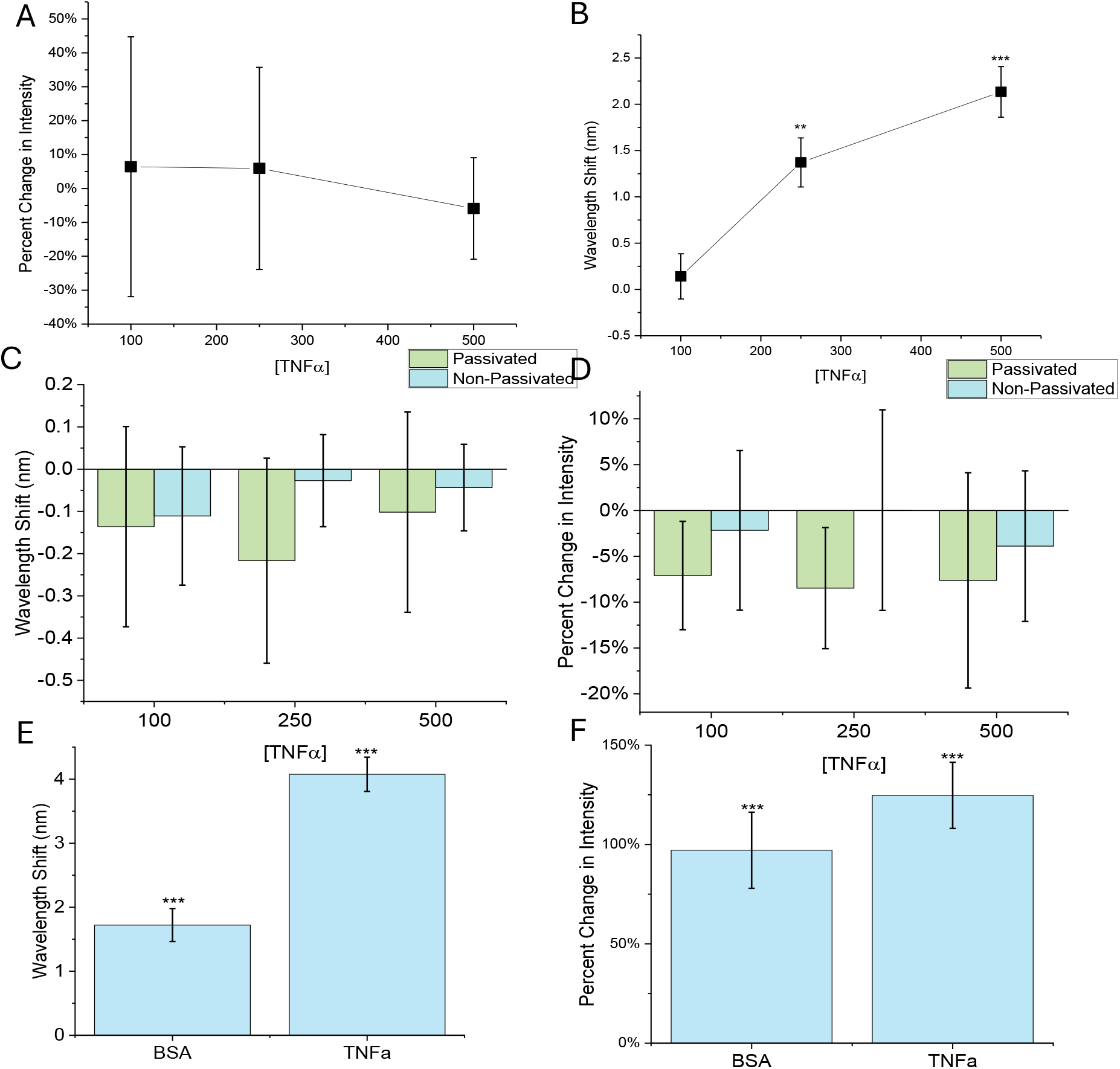
RNAapt-SWCNT sensor function in response to TNF-α. A) Intensity change and B) wavelength shift of the (7,6) SWCNT as a function of TNF-α concentration in 1x PBS. C) Comparison of the wavelength shift and D) intensity change of the (7,6) after 180 minutes in the presence of TNF-α in 10% FBS conditions with and without PLK passivation. E) Wavelength shift and F) intensity change of the (7,6) SWCNT in the presence of 250 nM BSA and TNF-α after 180 minutes in 1x PBS. Mean represents average of triplicate. Error bars represent +/-standard deviation. T-test significance indicated by, **=p<0.01; ***=p<0.001.

#### Monoclonal Antibody (mAb) conjugated ssDNA-SWCNT

Antibodies have a general reputation as highly specific and selective towards their analyte of interest. We therefore developed a TNF-α antibody-based nanosensor based simple conjugation of the antibody to ssDNA encapsulating SWCNT. Initial screening of the sensor against 250 nM TNF-α did not result in any noticeable changes in wavelength shift or intensity in both buffer and serum environments (**Figure 1**). Despite passivating the sensor with PLK to block nonspecific interactions in serum environments, it did not facilitate TNF-α detection.

#### Polyclonal Antibody (pAb) conjugated ssDNA-SWCNT

As antibodies exhibit varying utility in various contexts, it is important to screen different antibodies against a given target.^10, 11, 33^ For this reason, we tested a TNF-α-specific polyclonal antibody conjugated to ssDNA encapsulating SWCNT. Initial screening of the sensor in 1x PBS upon exposure to 250 nM TNF-α exhibit a wavelength shift of 1 nm in all (*n,m*) species analyzed, which was statistically significant for the (7,5) and (7,6) (**Figure 1, Supplementary Figure 2**). When challenged in 10% FBS, the passivated pAb-SWCNT sensor did not produce the response seen under buffer conditions. For this reason, more optimization of the sensor is required, as well as further validation of other TNF-α specific antibodies.

### Perspectives and future work

In our exploration of various molecular recognition elements complexed with SWCNT to detect TNF-α, we compared our results with the standards established in existing literature. The physiologically relevant range for TNF-α levels in humans is around 29 pM (∼5 pg/mL) in healthy individuals and anywhere from 29 nM to 290 nM (∼500 pg/mL to 5 ng/mL) in inflammatory disease states, depending on the biofluid and disease.^41, 42^ Higher values in the ng/mL range have been reported for animal and cell models, although they are not typically observed in humans. Conventional methods for TNF-α detection often rely on antibodies for their sensitivity and specificity. ELISA enables TNF-α quantification in the low nanomolar range, but is the process is time-consuming and prone to cross-reactivity.^43^ Other antibody-based techniques such as Western blot, flow cytometry, and immunohistochemistry are labor-intensive and require costly equipment. Electrochemical and optical sensors seek to provide alternatives for TNF-α quantification through methods which are highly sensitive, specific, inexpensive, rapid, and may even enable multiplexing.

Electrochemical impedance spectroscopy has been used in several instances to detect TNF-α in a variety of conditions, such as serum and saliva samples. Limits of detection are as low as 11.7 pM and multiplexing with other cytokines has also been reported.^44-50^ Antibodies are also used as the biorecognition element in these studies, typically immobilized on an electrode which conveys the TNF-α binding event through a change in electron transfer resistance. The polyclonal antibody tested in the present study also enabled detection of TNF-α when immobilized on SWCNT, though it was not functional in serum, suggesting further evaluation and optimization of this sensor design is needed for utility in early disease diagnostics. Cyclic voltammetry is another technique often used in electrochemical sensors, which uses a redox-active mediator or label to generate a current. Methylene blue-modified versions of VR11 and RNAapt have been used in this instance to detect 100 – 575 nM TNF-α.^34, 51^ Though these detection limits are higher than those of the antibody-based electrochemical sensors, they are still within the physiologically relevant range for TNF-α. In general, electrochemical sensors offer low costs and rapid response times but may suffer from interference issues and have limited utility *in vivo*.

Although no optical SWCNT sensors for TNF-α have been reported, other types of optical sensors such as surface plasmon resonance (SPR), fluorescence resonance energy transfer (FRET), and fiber optics have been published. Optical sensors hold many advantages, such as multiplexing ability and potential for spatial resolution when used *in vitro* or *in vivo*. Anti-TNF-α antibodies enabled detection of the cytokine as low as 1 nM with SPR and 16 pM with fiber optics.^52, 53^ VR11 was used in a FRET sensor made with quantum dots and gold nanoparticles, which reported a LOD of 98 nM.^35^ In the present study, SWCNT functionalized with VR11 exhibited optical detection as low as 1 nM.

Further work is needed to improve the LOD of the sensor constructs developed in this study. We explored the effects of thermal and ion-induced refolding of DNA on SWCNT, addition of a quencher dye, and use of flanking sequences to improve the function of VR11 as a recognition element for SWCNT sensors. Although none of these methods improved the sensitivity of our sensors in this study, further modifications of the aptamer sequences may still be explored, such as incorporating anchor sequences and chemical spacers or redox-active groups.^36^ Recent studies have also utilized solvatochromic dyes to improve the sensitivity of SWCNT-DNA sensors towards their target analytes.^54^ Furthermore, the optical properties of the SWCNT themselves can be improved through separation of chiral species, which eliminates spectral overlap and potentiates highly sensitive SWCNT constructs.

Most sensors also demonstrated limited specificity to TNF-α in this study compared to BSA. Surface passivation was used on the antibody-based sensors in this study, but future work could include a screening of various passivation agents to determine which can inhibit non-specific adsorption of biological proteins to the SWCNT surface.^28^

## Conclusions

This study presented methods to screen rationally designed SWCNT sensors for the detection of TNF-α. We prioritized known aptamers and variants thereof, with antibody-based sensors as comparators. We then evaluated the sensitivity and specificity of various sensor constructs and explored ways to improve sensor function. Several sensor constructs exhibited TNF-α sensitivity in buffer, though we found little success in detecting the cytokine in serum. We did find that several heat-cool cycles for 40Apt also induced sensitivity where previously there was none, while the RNA aptamer was able to discriminate TNF-α from BSA relatively well. We also found that addition of a Black Hole Quencher to the VR11 aptamer enhanced discrimination of the cytokine from BSA, while exacerbating its magnitude of change. This VR11-BHQ aptamer was, in fact, the only sensor that demonstrated function in serum conditions, which was facilitated by PLK passivation. Comparatively, the polyclonal antibody we used demonstrated some success in detecting TNF-α, however this was diminished in serum.

Our own prior experience in developing molecular recognition element-functionalized SWCNT is that it typically requires screening of several iterations of a single or multiple recognition elements to obtain a functional sensor.^7-10 11, 12^ This could in part be due to the inherent variability in commercially-produced antibodies^55-58^, or even that those that work in one assay may not work in other assays. It may also be because both DNA and RNA aptamers typically work best, or only, in the buffer and temperature conditions in which they were selected.^59-61^ Then, even assuming the best for the recognition element, the interfacial conditions in SWCNT-based sensors are substantially different than other types of molecular assays or diagnostic sensors. These screening efforts often assess five or ten different binding elements, though the full set trial-and-error is rarely disseminated. In this work, we aimed to provide a rational framework for screening recognition elements for rationally-designed SWCNT sensors. We began with an initial screen of sensor constructs based on literature studies and sought to improve upon promising leads through several methods. We are confident that at least one sensor construct, likely with the VR11-BHQ variant, will have utility for in vitro and in vivo assays, or personal diagnostics, against the important pro-inflammatory cytokine TNF-α.

## Acknowledgements

The authors wish to acknowledge all members of the Williams Lab for discussion and feedback. This work was supported by NIH R35GM142833, The City College of New York Grove School of Engineering, Stony Brook University Department of Medicine, and the SUNY Empire Innovation Program Award #250010 (R. Williams). A. Ryan and A. Israel were supported by a G-RISE Ph.D. traineeship from the National Institutes of Health (T32GM136499).

## References

(1) Turner, M. D.; Nedjai, B.; Hurst, T.; Pennington, D. J. Cytokines and chemokines: At the crossroads of cell signalling and inflammatory disease. Biochimica et Biophysica Acta (BBA) - Molecular Cell Research 2014, 1843 (11), 2563–2582.

(2) Kany, S.; Vollrath, J. T.; Relja, B. Cytokines in Inflammatory Disease. Int J Mol Sci 2019, 20 (23), 6008.

(3) Bradley, J. TNF-mediated inflammatory disease. The Journal of Pathology 2008, 214 (2), 149–160.

(4) Jang, D. I.; Lee, A. H.; Shin, H. Y.; Song, H. R.; Park, J. H.; Kang, T. B.; et al. The Role of Tumor Necrosis Factor Alpha (TNF-α) in Autoimmune Disease and Current TNF-α Inhibitors in Therapeutics. Int J Mol Sci 2021, 22 (5).

(5) Ackermann, J.; Metternich, J. T.; Herbertz, S.; Kruss, S. Biosensing with Fluorescent Carbon Nanotubes. Angewandte Chemie International Edition 2022, 61 (18), e202112372.

(6) Cohen, Z.; Williams, R. M. Single-walled carbon nanotubes as optical transducers for nanobiosensors in vivo. ACS nano 2024, 18 (52), 35164–35181.

(7) Ryan, A. K.; Rahman, S.; Williams, R. M. Optical Aptamer-Based Cytokine Nanosensor Detects Macrophage Activation by Bacterial Toxins. ACS Sensors 2024, 9 (7), 3697–3706.

(8) Zanetti, J. K.; Stefoni, M. C.; Ferraz, C.; Ryan, A.; Israel, A.; Williams, R. M. A near-infrared fluorescent aptananosensor enables selective detection of the stress hormone cortisol in artificial cerebrospinal fluid. Sensors & Diagnostics 2025, 4 (12), 1103–1113.

(9) Williams, R. M.; Lee, C.; Galassi, T. V.; Harvey, J. D.; Leicher, R.; Sirenko, M.; et al. Noninvasive ovarian cancer biomarker detection via an optical nanosensor implant. Science Advances 2018, 4 (4), eaaq1090.

(10) Gaikwad, P.; Rahman, N.; Parikh, R.; Crespo, J.; Cohen, Z.; Williams, R. M. Optical Nanosensor Passivation Enables Highly Sensitive Detection of the Inflammatory Cytokine Interleukin-6. ACS Applied Materials & Interfaces 2024, 16 (21), 27102–27113.

(11) Williams, R. M.; Lee, C.; Heller, D. A. A Fluorescent Carbon Nanotube Sensor Detects the Metastatic Prostate Cancer Biomarker uPA. ACS Sensors 2018, 3 (9), 1838–1845.

(12) Gaikwad, P. V.; Rahman, N.; Ghosh, P.; Ng, D. L.; Williams, R. M. Detection of Estrogen Receptor Status in Breast Cancer Cytology Samples by an Optical Nanosensor. Advanced NanoBiomed Research 2025, 5 (1), 2400099.

(13) Zheng, M.; Jagota, A.; Semke, E. D.; Diner, B. A.; McLean, R. S.; Lustig, S. R.; et al. DNA-assisted dispersion and separation of carbon nanotubes. Nat Mater 2003, 2 (5), 338–342.

(14) Orava, E. W.; Jarvik, N.; Shek, Y. L.; Sidhu, S. S.; Gariépy, J. A Short DNA Aptamer That Recognizes TNFα and Blocks Its Activity in Vitro. ACS Chemical Biology 2013, 8 (1), 170–178.

(15) Bruno, J. G.; Carrillo, M. P.; Phillips, T.; Andrews, C. J. A Novel Screening Method for Competitive FRET-Aptamers Applied to E. coli Assay Development. Journal of Fluorescence 2010, 20 (6), 1211–1223.

(16) Hwang, J. Y.; Kim, S. T.; Han, H.-S.; Kim, K.; Han, J. S. Optical Aptamer Probes of Fluorescent Imaging to Rapid Monitoring of Circulating Tumor Cell. Sensors 2016, 16 (11), 1909.

(17) Yang, X.; Han, Q.; Zhang, Y.; Wu, J.; Tang, X.; Dong, C.; et al. Determination of free tryptophan in serum with aptamer—Comparison of two aptasensors. Talanta 2015, 131, 672–677.

(18) Moutsiopoulou, A.; Broyles, D.; Dikici, E.; Daunert, S.; Deo, S. K. Molecular Aptamer Beacons and Their Applications in Sensing, Imaging, and Diagnostics. Small 2019, 15 (35), 1902248.

(19) Yu, H.; Zhao, Q. Aptamer Molecular Beacon Sensor for Rapid and Sensitive Detection of Ochratoxin A. Molecules 2022, 27 (23).

(20) Li, Y.; Sun, L.; Zhao, Q. Development of aptamer fluorescent switch assay for aflatoxin B1 by using fluorescein-labeled aptamer and black hole quencher 1-labeled complementary DNA. Anal Bioanal Chem 2018, 410 (24), 6269–6277.

(21) Lai, W. Y.; Wang, J. W.; Huang, B. T.; Lin, E. P.; Yang, P. C. A Novel TNF-α-Targeting Aptamer for TNF-α-Mediated Acute Lung Injury and Acute Liver Failure. Theranostics 2019, 9 (6), 1741–1751.

(22) Liu, Y.; Zhou, Q.; Revzin, A. An aptasensor for electrochemical detection of tumor necrosis factor in human blood. Analyst 2013, 138 (15), 4321–4326.

(23) Landry, M. P.; Vuković, L.; Kruss, S.; Bisker, G.; Landry, A. M.; Islam, S.; et al. Comparative Dynamics and Sequence Dependence of DNA and RNA Binding to Single Walled Carbon Nanotubes. The Journal of Physical Chemistry C 2015, 119 (18), 10048–10058.

(24) Mann, F. A.; Herrmann, N.; Meyer, D.; Kruss, S. Tuning Selectivity of Fluorescent Carbon Nanotube-Based Neurotransmitter Sensors. Sensors (Basel) 2017, 17 (7).

(25) Lew, T. T. S.; Park, M.; Cui, J.; Strano, M. S. Plant Nanobionic Sensors for Arsenic Detection. Advanced Materials 2021, 33 (1), 2005683.

(26) Kruss, S.; Salem, D. P.; Vuković, L.; Lima, B.; Vander Ende, E.; Boyden, E. S.; et al. High-resolution imaging of cellular dopamine efflux using a fluorescent nanosensor array. Proceedings of the National Academy of Sciences 2017, 114 (8), 1789–1794.

(27) Giraldo, J. P.; Landry, M. P.; Kwak, S.-Y.; Jain, R. M.; Wong, M. H.; Iverson, N. M.; et al. A Ratiometric Sensor Using Single Chirality Near-Infrared Fluorescent Carbon Nanotubes: Application to In Vivo Monitoring. Small 2015, 11 (32), 3973–3984.

(28) Gaikwad, P.; Rahman, N.; Parikh, R.; Crespo, J.; Cohen, Z.; Williams, R. M. Optical Nanosensor Passivation Enables Highly Sensitive Detection of the Inflammatory Cytokine Interleukin-6. ACS Appl Mater Interfaces 2024, 16 (21), 27102–27113.

(29) Lee, K.; Nojoomi, A.; Jeon, J.; Lee, C. Y.; Yum, K. Near-Infrared Fluorescence Modulation of Refolded DNA Aptamer-Functionalized Single-Walled Carbon Nanotubes for Optical Sensing. ACS Applied Nano Materials 2018, 1 (9), 5327–5336.

(30) Weisman, R. B.; Bachilo, S. M. Dependence of Optical Transition Energies on Structure for Single-Walled Carbon Nanotubes in Aqueous Suspension: An Empirical Kataura Plot. Nano Letters 2003, 3 (9), 1235–1238.

(31) Bachilo, S. M.; Strano, M. S.; Kittrell, C.; Hauge, R. H.; Smalley, R. E.; Weisman, R. B. Structure-Assigned Optical Spectra of Single-Walled Carbon Nanotubes. Science 2002, 298 (5602), 2361–2366.

(32) Gaikwad, P.; Rahman, N.; Parikh, R.; Crespo, J.; Cohen, Z.; Williams, R. Optical nanosensor passivation enables highly sensitive detection of the inflammatory cytokine IL-6. bioRxiv 2023, 2023.2005.2010.540217.

(33) Williams, R. M.; Lee, C.; Galassi, T. V.; Harvey, J. D.; Leicher, R.; Sirenko, M.; et al. Noninvasive ovarian cancer biomarker detection via an optical nanosensor implant. Sci Adv 2018, 4 (4), eaaq1090.

(34) Mayer, M. D.; Lai, R. Y. Effects of redox label location on the performance of an electrochemical aptamer-based tumor necrosis factor-alpha sensor. Talanta 2018, 189, 585–591.

(35) Ghosh, S.; Datta, D.; Chaudhry, S.; Dutta, M.; Stroscio, M. A. Rapid Detection of Tumor Necrosis Factor-Alpha Using Quantum Dot-Based Optical Aptasensor. IEEE Transactions on NanoBioscience 2018, 17 (4), 417–423.

(36) Landry, M. P.; Ando, H.; Chen, A. Y.; Cao, J.; Kottadiel, V. I.; Chio, L.; et al. Single-molecule detection of protein efflux from microorganisms using fluorescent single-walled carbon nanotube sensor arrays. Nature Nanotechnology 2017, 12 (4), 368–377.

(37) Harvey, J. D.; Jena, P. V.; Baker, H. A.; Zerze, G. H.; Williams, R. M.; Galassi, T. V.; et al. A carbon nanotube reporter of microRNA hybridization events in vivo. Nat Biomed Eng 2017, 1 (4), 1–11.

(38) Li, Y.; Sun, L.; Zhao, Q. Development of aptamer fluorescent switch assay for aflatoxin B1 by using fluorescein-labeled aptamer and black hole quencher 1-labeled complementary DNA. Analytical and Bioanalytical Chemistry 2018, 410 (24), 6269–6277.

(39) Arrigo, R.; Teresi, R.; Gambarotti, C.; Parisi, F.; Lazzara, G.; Dintcheva, N. T. Sonication-Induced Modification of Carbon Nanotubes: Effect on the Rheological and Thermo-Oxidative Behaviour of Polymer-Based Nanocomposites. Materials (Basel) 2018, 11 (3).

(40) Fan, R.; Tao, X.; Zhai, X.; Zhu, Y.; Li, Y.; Chen, Y.; et al. Application of aptamer-drug delivery system in the therapy of breast cancer. Biomedicine & Pharmacotherapy 2023, 161, 114444.

(41) Kassasseya, C.; Torsin, L. I.; Musset, C.; Benhamou, M.; Chaudry, I. H.; Cavaillon, J.-M.; et al. Divergent effects of tumor necrosis factor (TNF) in sepsis: a meta-analysis of experimental studies. Critical Care 2024, 28 (1), 293.

(42) Gharamti, A. A.; Samara, O.; Monzon, A.; Montalbano, G.; Scherger, S.; DeSanto, K.; et al. Proinflammatory Cytokine Levels in Sepsis and in Health and TNFα Association with Sepsis Mortality and patient characteristics: a Systematic Review and Meta-analysis. medRxiv 2021, 2021.2012.2013.21267720.

(43) Chiswick, E. L.; Duffy, E.; Japp, B.; Remick, D. Detection and quantification of cytokines and other biomarkers. Methods Mol Biol 2012, 844, 15–30.

(44) La Belle, J. T.; Demirok, U. K.; Patel, D. R.; Cook, C. B. Development of a novel single sensor multiplexed marker assay. Analyst 2011, 136 (7), 1496–1501, 10.1039/C0AN00923G.

(45) Sri, S.; Chauhan, D.; Lakshmi, G. B. V. S.; Thakar, A.; Solanki, P. R. MoS2 nanoflower based electrochemical biosensor for TNF alpha detection in cancer patients. Electrochimica Acta 2022, 405, 139736.

(46) Bhatti, F.; Xiao, D.; Jebagu, T.; Huang, X.; Witherspoon, E.; Dong, P.; et al. Semiconductive biocomposites enabled portable and interchangeable sensor for early osteoarthritis joint inflammation detection. Advanced Composites and Hybrid Materials 2023, 6 (1), 33.

(47) Sánchez-Tirado, E.; Salvo, C.; González-Cortés, A.; Yáñez-Sedeño, P.; Langa, F.; Pingarrón, J. M. Electrochemical immunosensor for simultaneous determination of interleukin-1 beta and tumor necrosis factor alpha in serum and saliva using dual screen printed electrodes modified with functionalized double–walled carbon nanotubes. Analytica Chimica Acta 2017, 959, 66–73.

(48) Pui, T. S.; Kongsuphol, P.; Arya, S. K.; Bansal, T. Detection of tumor necrosis factor (TNF-α) in cell culture medium with label free electrochemical impedance spectroscopy. Sensors and Actuators B: Chemical 2013, 181, 494–500.

(49) Say, R.; Özkütük, E. B.; Ünlüer, Ö. B.; Uğurağ, D.; Ersöz, A. Nano anti-tumor necrosis factor-alpha based potentiometric sensor for tumor necrosis factor-alpha detection. Sensors and Actuators B: Chemical 2015, 209, 864–869.

(50) Wei, H.; Ni, S.; Cao, C.; Yang, G.; Liu, G. Graphene Oxide Signal Reporter Based Multifunctional Immunosensing Platform for Amperometric Profiling of Multiple Cytokines in Serum. ACS Sens. 2018, 3 (8), 1553–1561.

(51) Liu, Y.; Kwa, T.; Revzin, A. Simultaneous detection of cell-secreted TNF-α and IFN-γ using micropatterned aptamer-modified electrodes. Biomaterials 2012, 33 (30), 7347–7355.

(52) Predabon, S.; Buzzetti, P. H.; Visentainer, J. E.; Visentainer, J.; Radovanovic, E.; Monteiro, J.; et al. Detection of tumor necrosis factor-alpha cytokine from the blood serum of a rat infected with Pb18 by a gold nanohole array-based plasmonic biosensor. Journal of Nanophotonics 2020, 14 (3), 036004.

(53) Cao, H.; Liu, J.; Wei, D.; Liu, B.; Hu, Y.; Liu, J.; et al. Ultrahigh sensitivity fiber laser integrated biosensor for TNF-α detection in human blood sample. Sensors and Actuators B: Chemical 2025, 422, 136561.

(54) Ma, C.; Kistwal, T.; Hill, B. F.; Neutsch, K.; Kruss, S. Solvatochromic Dyes Increase the Sensitivity of Nanosensors. J. Phys. Chem. C 2025, 129 (3), 1824–1830.

(55) Laflamme, C.; McKeever, P. M.; Kumar, R.; Schwartz, J.; Kolahdouzan, M.; Chen, C. X.; et al. Implementation of an antibody characterization procedure and application to the major ALS/FTD disease gene C9ORF72. Elife 2019, 8, e48363.

(56) Kwon, D. The antibodies don’t work! The race to rid labs of molecules that ruin experiments. Nature 2024, 635 (8037), 26–28.

(57) A Kahn, R.; Virk, H.; Laflamme, C.; W Houston, D.; Polinski, N. K.; Meijers, R.; et al. Science Forum: Antibody characterization is critical to enhance reproducibility in biomedical research. 2024.

(58) Ayoubi, R.; Ryan, J.; Biddle, M. S.; Alshafie, W.; Fotouhi, M.; Bolivar, S. G.; et al. Scaling of an antibody validation procedure enables quantification of antibody performance in major research applications. Elife 2023, 12, RP91645.

(59) Miller, A. A.; Rao, A. S.; Nelakanti, S. R.; Kujalowicz, C.; Shi, T.; Rodriguez, T.; et al. Systematic review of aptamer sequence reporting in the literature reveals widespread unexplained sequence alterations. Analytical chemistry 2022, 94 (22), 7731–7737.

(60) Arroyo-Currás, N.; Dauphin-Ducharme, P.; Scida, K.; Chávez, J. L. From the beaker to the body: translational challenges for electrochemical, aptamer-based sensors. Analytical Methods 2020, 12 (10), 1288–1310.

(61) Arroyo-Currás, N. Aptamers Can Be Effective Affinity Receptors for Biosensing. ECS Sensors Plus 2024, 3 (3), 030001.

