## Supplementary Figures for "Screening Molecular Recognition Element-Based SWCNT Optical Sensors for the Inflammatory Cytokine TNF-α"

### Supporting Figures

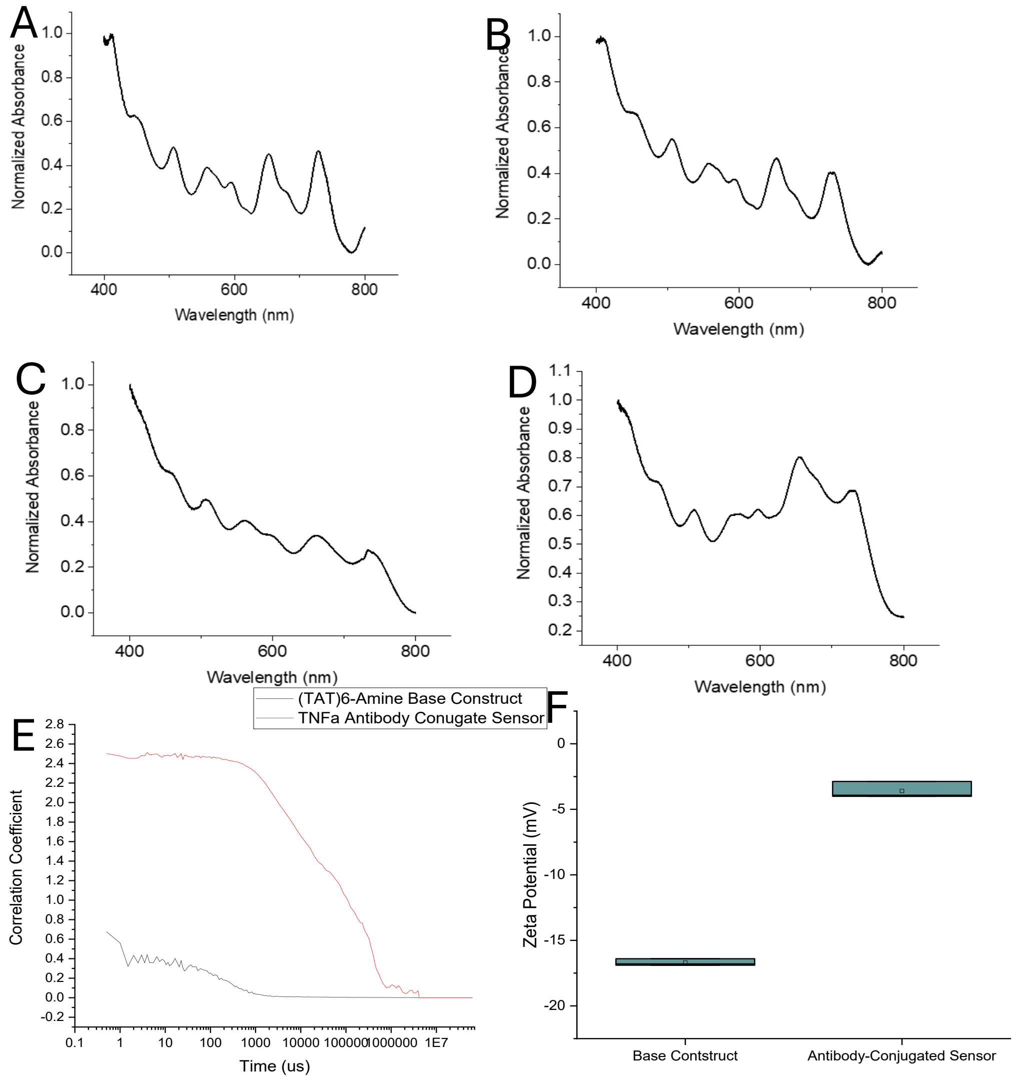


**Supplementary Figure 1. Absorbance and DLS of sensor constructs.** A) Absorbance spectra of VR11-SWCNT, B) $(GT)_{15}$ C) SWCNT-RNAapt D) SWCNT-VR11-BHQ. E) Dynamic light scattering (DLS) of the $(TAT)_6$-$NH_2$-SWCNT before and after the conjugation of the monoclonal TNF-α antibody. F) ζ-potential of the $(TAT)_6$-$NH_2$-SWCNT before and after the conjugation of the monoclonal TNF-α antibody.

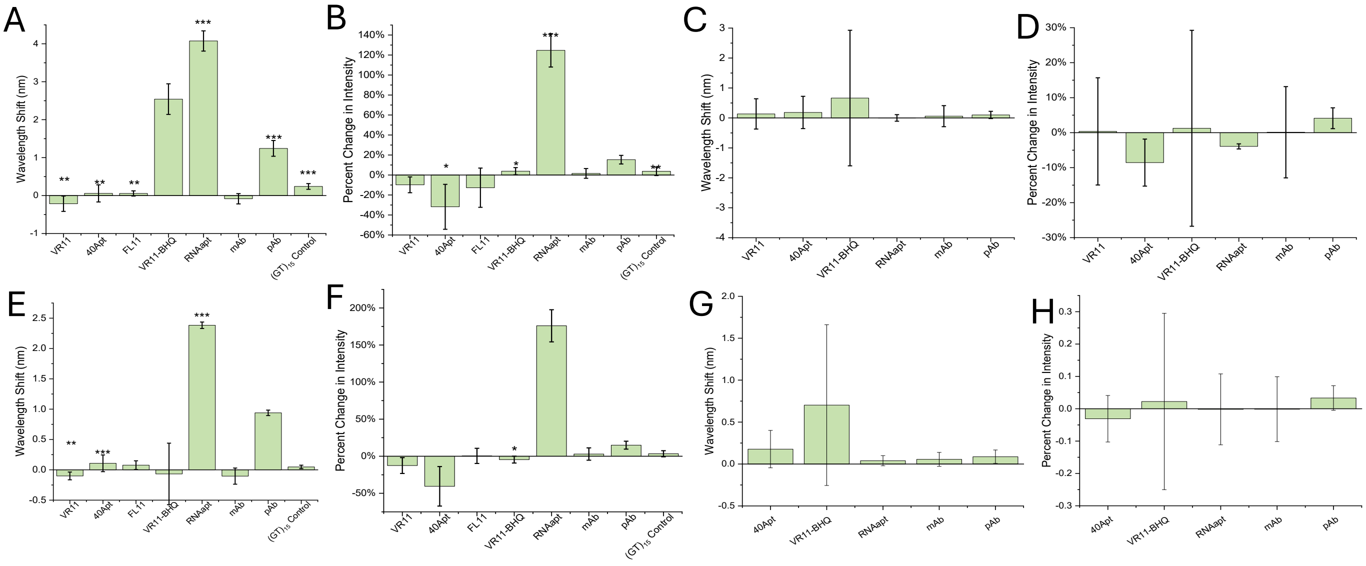


**Supplementary Figure 2. (7,6) and (9,4) SWCNT intensity and wavelength changes.** A) Wavelength shift of the (7,6) species of all sensor constructs after three hours of incubation with 250 nM TNF-α protein in 1X PBS. B) Changes in the (7,6) emission intensity. C) Wavelength shift of the (7,6) chirality of all sensor constructs after three hours of incubation with 250 nM TNF-α protein in 10% FBS. D) Changes in the (7,6) emission intensity E) Wavelength shift of the (9,4) species of all sensor constructs after three hours of incubation with 250 nM TNF-α protein in 1X PBS. F) Changes in the (9,4) emission intensity. G) Wavelength shift of the (9,4) species of all sensor constructs after three hours of incubation with 250 nM TNF-α protein in 10% FBS. H) Changes in the (9,4) emission intensity. Error bars represent +/- standard deviation. T-test significance indicated by* =$p<0.05$, **=$p<0.01$, ***=$p<0.001$

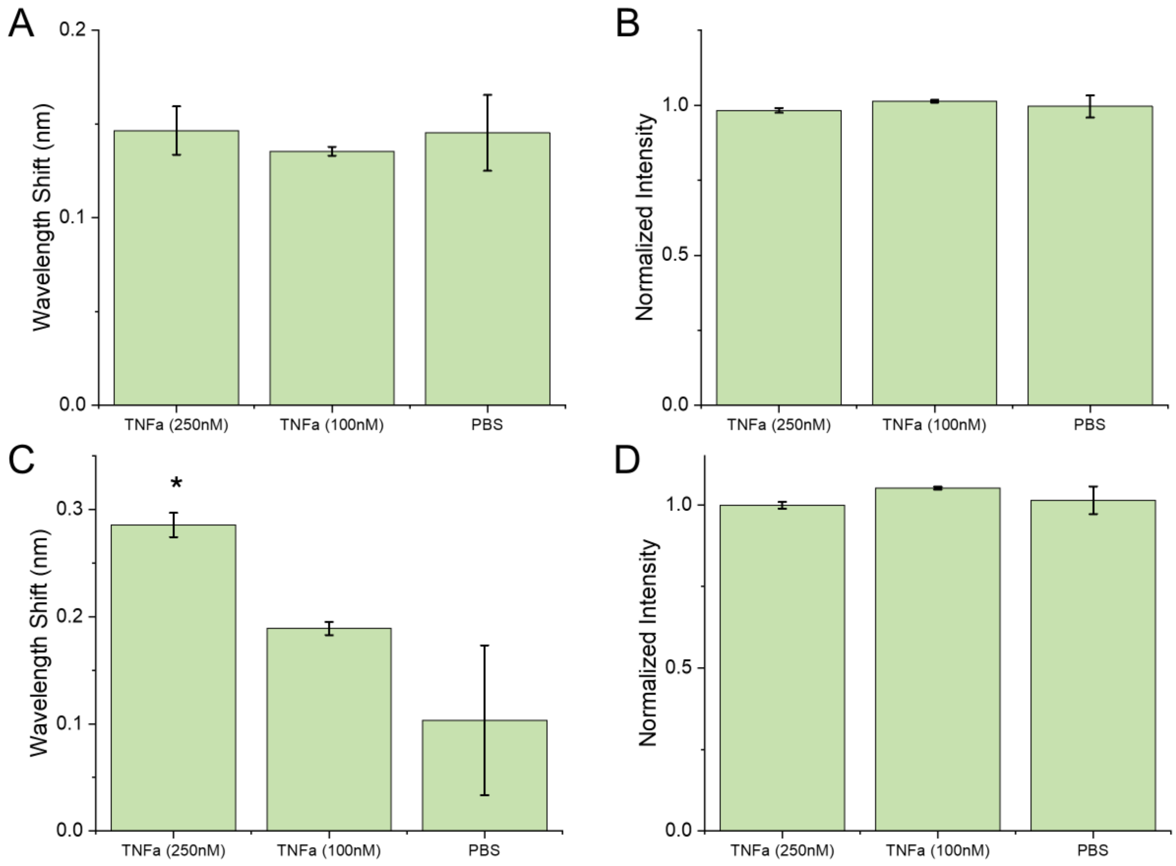


**Supplementary Figure 3. Changes in $(GT)_{15}$-SWCNT control.** A) Shifts in center wavelength and B) intensity of the (7,5) SWCNT after 180 minute TNF-α incubation in 1x PBS. C) Shifts in center wavelength and B) intensity of the (7,6) SWCNT after 180 minutes incubation in 1x PBS. Mean represents average of triplicate. Error bars represent +/- standard deviation. T-test significance indicated by* =$p<0.05$, **=$p<0.01$, ***=$p<0.001$.

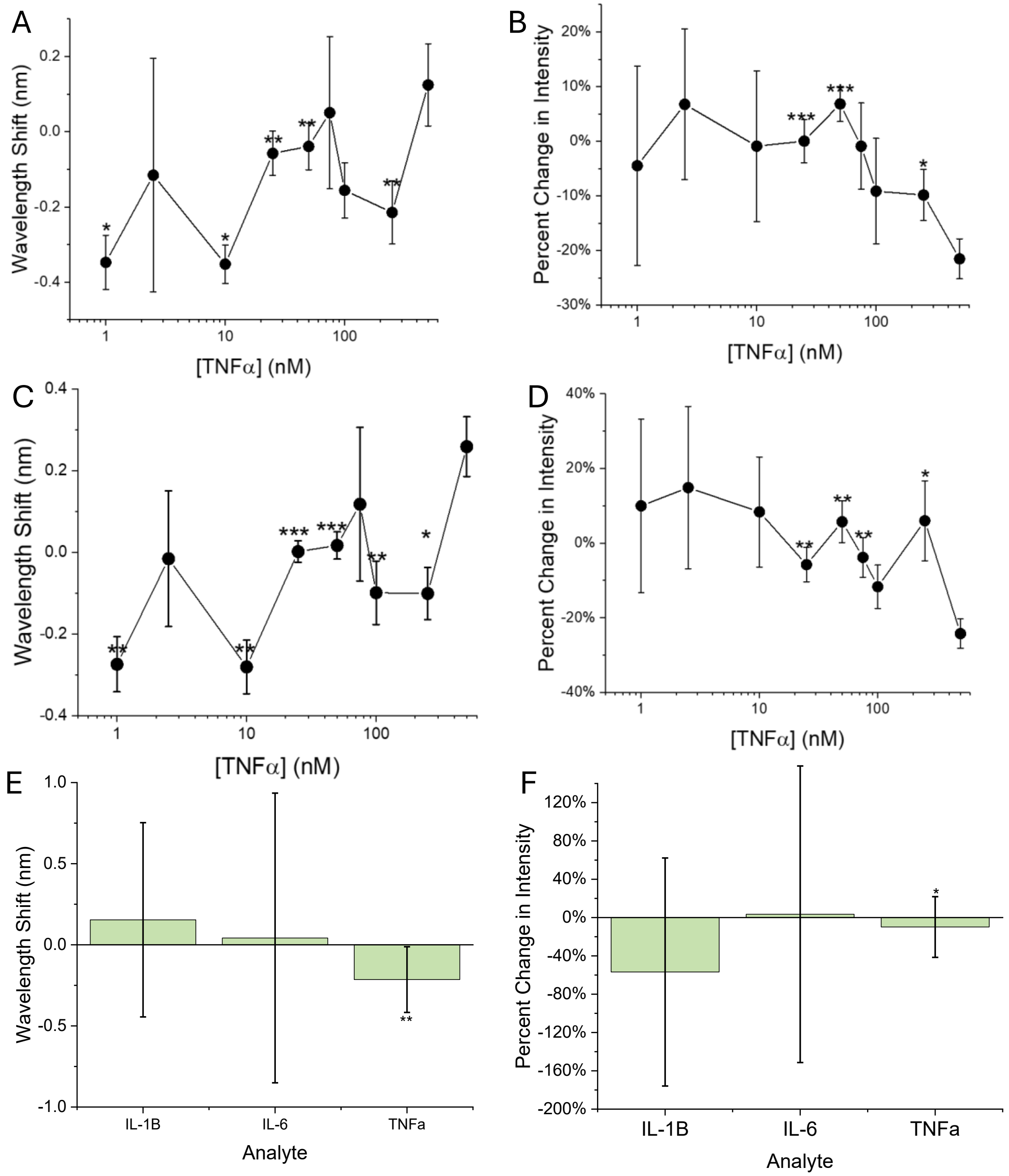


**Supplementary Figure 4. VR11-SWCNT sensor assessment**. A) Shifts in center wavelength of the (7,6) SWCNT after TNF-α after 180 minute incubation performed in 1x PBS and B) modulations of (7,6) emission intensity. C) Shifts in center wavelength of the (9,4) SWCNT at varying concentrations after 180 minutes incubation performed in 1x PBS and D) modulations of (9,4) emission intensity. E) Wavelength shift and F) intensity change of the (7,6) after incubation with 250 nM IL-1β, IL-6, and TNF-α. Mean represents average of triplicate. Error bars represent +/- standard deviation. T-test significance indicated by* $=p<0.05$, **$=p<0.01$, ***$=p<0.001$.

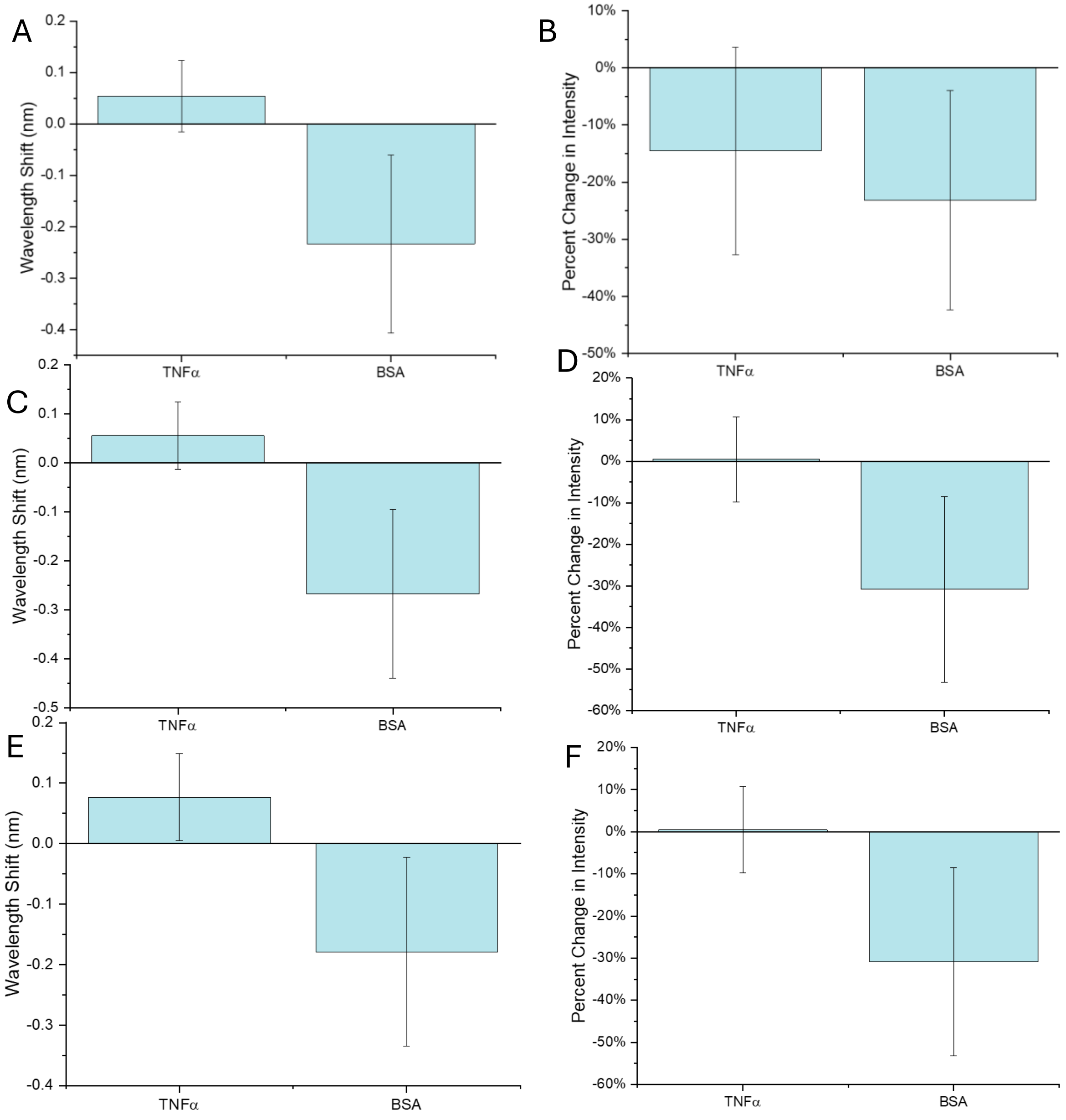


**Supplementary Figure 5. FL11-SWCNT sensor assessment**. A) Shift in center wavelength and B) change in intensity of the (7,5) after 180 minutes incubation with BSA and TNF-α in 1x PBS. C) Shift in center wavelength and D) change in intensity of the (7,6) after 180 minutes incubation with BSA and TNF-α in 1x PBS. E) Shift in center wavelength and F) intensity of the (9,4) after 180 minutes incubation with BSA and TNF-α in 1x PBS.

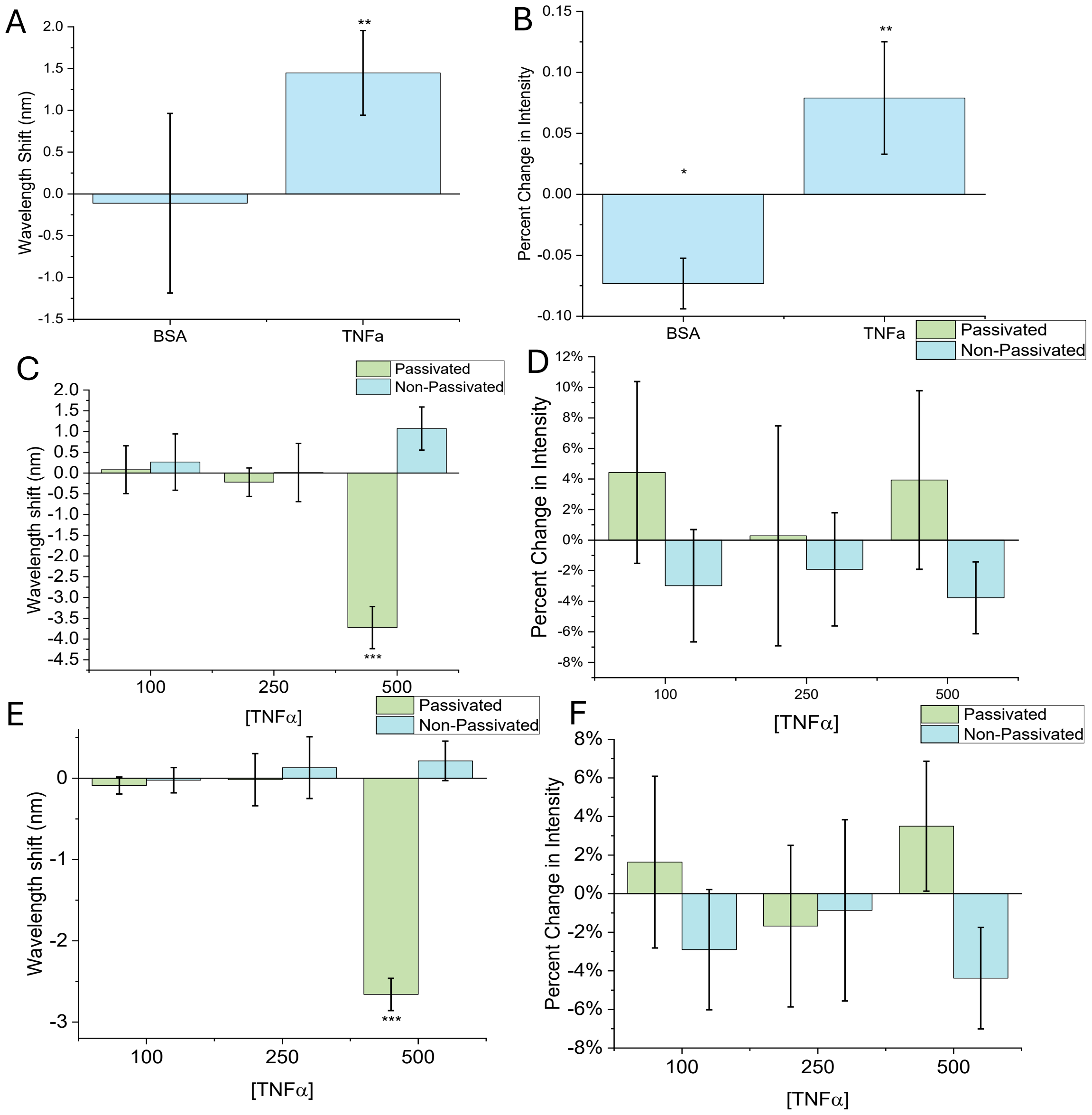


**Supplementary Figure 6. VR11-BHQ-SWCNT sensor assessment.** A) Shift in center wavelength of the (9,4) after 180 minutes incubation with BSA and TNF-α in 1x PBS. B) Modulations in the (9,4) emission intensity after 180 minutes incubation with BSA and TNF-α in 1x PBS. C) Comparison of the wavelength shift of the (7,5) after 180 minutes in the presence of TNF-α of the VR11-BHQ-SWCNT construct in 10% FBS conditions with and without PLK passivation. D) Comparison of intensity modulations in the (7,5) after 180 minutes in the presence of TNF-α of the RNAapt-SWCNT construct in 10% FBS conditions with and without PLK passivation. E) Comparison of the wavelength shift of the (9,4) after 180 minutes in the presence of TNF-α of the VR11-BHQ-SWCNT construct in 10% FBS conditions with and without PLK passivation. F) Comparison of intensity modulations in the (9,4) after 180 minutes in the presence of TNF-α of the VR11-BHQ-SWCNT construct in 10% FBS conditions with and without PLK passivation.

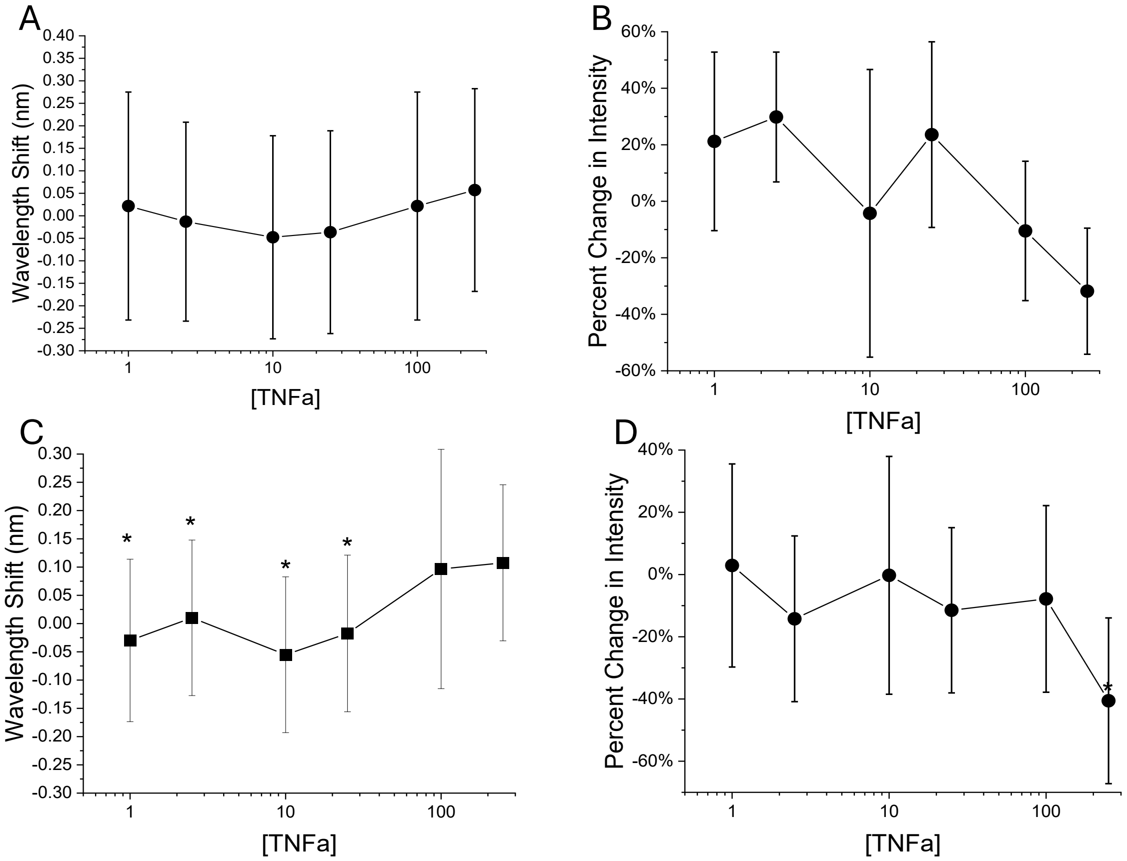


**Supplementary Figure 7. 40Apt-SWCNT concentration-dependent response to TNF-α**. A) Shifts in center wavelength of the (7,6) at each concentration after 180 minute incubation performed in 1xPBS B) Modulations of (7,6) emission intensity. C) Shifts in center wavelength of the (9,4) at each concentration after 180 minutes incubation in 1x PBS D) Modulations of (9,4) emission intensity Mean represents average of triplicate. Error bars represent +/- standard deviation. T-test significance indicated by* =p<0.05, **=p<0.01, ***=p<0.001

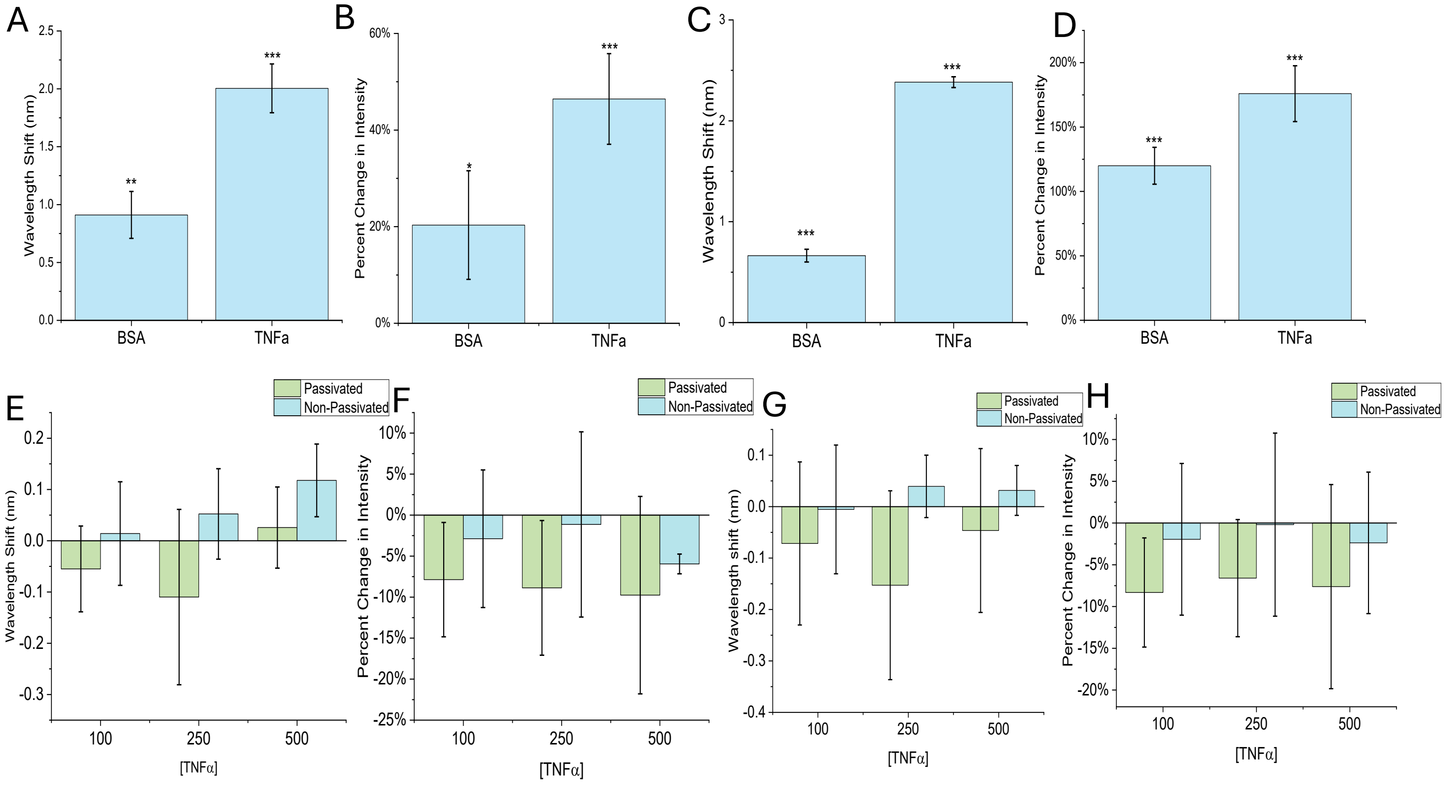


**Supplementary Figure 8. RNAapt-SWCNT sensor assessment.** A) Shift in center wavelength of the (7,5) after 180 minutes incubation with BSA and TNF-α in 1x PBS. B) Modulations in the (7,5) emission intensity after 180 minutes incubation with BSA and TNF-α in 1x PBS. C) Shift in center wavelength of the (9,4) after 180 minutes incubation with BSA and TNF-α in 1x PBS. D) Modulations in the (9,4) emission intensity after 180 minutes incubation with BSA and TNF-α in 1x PBS. E) Comparison of the wavelength shift of the (7,5) after 180 minutes in the presence of TNF-α of the RNAapt-SWCNT construct in 10% FBS conditions with and without PLK passivation. F) Comparison of intensity modulations in the (7,5) after 180 minutes in the presence of TNF-α of the RNAapt-SWCNT construct in 10% FBS conditions with and without PLK passivation. G) Comparison of the wavelength shift of the (9,4) after 180 minutes in the presence of TNF-α of the RNAapt-SWCNT construct in 10% FBS conditions with and without PLK passivation. H) Comparison of intensity modulations in the (9,4) after 180 minutes in the presence of TNF-α of the RNAapt-SWCNT construct in 10% FBS conditions with and without PLK passivation
